# Fragrance and color production by corona and perianth tissues of Iranian narcissus genotypes (*Narcissus tazetta* L.)

**DOI:** 10.1101/2023.10.21.563448

**Authors:** Azra Haghshenas, Abolfazl Jowkar, Mehrangiz Chehrazi, Ali Moghadam, Akbar Karami

## Abstract

Flower color, shape and scent are the most attracting factors for consumers in the floriculture industry. The most fragrant Iranian narcissi (*Narcissus tazetta* L.) grown in natural habitats are Shahla, Meskinak, and Porpar genotypes. The present study was conducted to evaluate the color, scent, and also their interaction separately in corona and perianth of eleven Iranian narcissus accessions, for a better understanding of the bio-physiological differences in these tissues. For this purpose, the volatile organic compounds (VOCs) were analyzed using Headspace GCMS (gas chromatography/mass spectrometry); and total carotenoid, color values, and tissue thickness were measured in both perianth and corona tissues. Sensory analysis for corona and perianth was also conducted to evaluate scent perception. Besides, the expression of genes related to scent and color in corona and perianth was evaluated. Moreover, epidermal cells of perianth and corona were assessed by polarized light and SEM microscopy. The two most abundant compounds in both tissues were E-β-ocimene and benzyl acetate, respectively; among which the first compound was higher in perianth, while the second one was higher in corona. Among identified VOCs, α-terpineol, acetic acid, 2-phenylethyl ester, α-pinene, β-myrcene, and limonene were detected in corona of all genotypes of *N. tazetta*; however, these compounds were not identified in perianth of every genotypes. In corona, the negative correlations between the measured values of E-β-ocimene and carotenoid and also, between the expression level of ocimene synthase and PSY (Phytoene synthase) with DXR (1-Deoxy-D-xylulose 5-phosphate reductoisomerase) suggested that there might be a competition between carotenoids and monoterpenes precursors in the MEP (methyl-D-erythritol phosphate) pathway. Volatile organic compound, color, scent emission, fresh weight and thickness of tissue were different in perianth and corona; while the surface coverage (with epidermal conical cells) were similar in these tissues. The findings of this research illustrated clearly for the first time that while both perianth and corona play important roles in scent production, corona has a more distinguished role in greater production of scent and color in Iranian narcissus flowers.

## 1. Introduction

Floral shape, color, and scent are visual and chemical signals that attract pollinators and secure plants’ reproductive and evolutionary success. These factors are also important aesthetic features in attracting consumers to the products of the floriculture industry (Anisworth, 2006). The shape and color of flower are two important characteristics that absorb the customer’s attention at the first glance. Often, evaluation of morphological factors is much easier than scent assessment. Until recently, before the invention of instruments and methods of analyzing volatile organic compounds (VOCs) which has greatly contributed to the research of flower scents, evaluations conducted on the flower color were much more frequent compared with the flower scent.

The Iranian narcissus (*Narcissus tazetta*) is native to West Asia and the Mediterranean region, and has been introduced into numerous countries (Kamenetsky and Okubo, 2012). It has several natural habitats, among which the largest ones are located in southern Iran. Shahla, Meskinak, Porpar, and Panje-gorbehi (i.e. cat-paws) are the common genotypes, among which the first three ones are the most fragrant. In addition to natural habitats, Shahla and Porpar are cultivated in farms or under trees of orchards for their good marketability. Meskinak has strong scent; however, it is not cultivated due to the short vase life and small size of flowers. Besides the aesthetic aspects of Iranian narcissus, in Persian traditional medicine and Unani medicine it is well-known that smelling fragrant *N. tazetta* flowers is beneficial for aromatherapy to cure cold (Aghili Khorasani, 2009; Kalam & Qayoom, 2020), epilepsy and cardiac problems, and to clear the obstruction in brain blood vessels (Qayoom, 2020).

Floral scent is created by a mixture of low molecular weight compounds, emitted from flowers. These compounds are comprised of three major groups: Terpenoids, phenylpropanoids/benzenoids and fatty acid derivatives. The main scent compounds of *N. tazetta* include monoterpenes such as E-β-ocimene (Ruíz-Ramón et al., 2014; Zarifikhosroshahi et al., 2021; and He et al., 2020), Eucalyptol (Ruíz-Ramón et al., 2014), linalool (Zarifikhosroshahi et al., 2021) and benzenoid (benzyl acetate), among which E-β-ocimene has the most percentage (Ruíz-Ramón et al., 2014; Zarifikhosroshahi et al., 2021). A few biochemical pathways synthesize the flower volatiles (Anisworth, 2006). Monoterpenes are synthesized through MEP (methyl-D-erythritol phosphate) pathway, where E-β-ocimene is produced by the E-β-ocimene synthase enzyme. The responsible gene for E-β-ocimene synthase belongs to the TPS (terpene synthase) gene family which could synthetize one or more terpenoids (Farré-Armengol et al., 2017; and Dudareva et al., 2003).

The orange color of narcissus flower is due to the carotenoid pigment. Li et al. (2015) identified ten carotenoids and seventeen flavonols in narcissus cultivars with orange and pink colors. Also, Ren et al. (2017) reported that in narcissus, carotenoid metabolic pathway is the main contributor to the flower color. Quantity of carotenoid in flowers depends on the equilibrium between biosynthesis and degradation of this compound. In the carotenoid biosynthesis, product of PSY (phytoene synthase) is the first and important rate-limiting plastidial enzyme which converts two molecules of geranylgeranyl diphosphate (GGPP) to phytoene. In carotenoid biodegradation, cleavage deoxygenases (CCDs) and 9-*cis*-Epoxycarotenoid dioxygenases (NCEDs) degrade carotenoids to other compounds; however, up to now, no NCED gene has been identified in narcissus species (Zhang et al. 2022). Carotenoids and monoterpenes are both produced by the MEP pathway in plastids. DXS (1-deoxy-D-xylulose-5-phosphate synthase; Xu et al., 2016) and DXR (1-Deoxy-D-xylulose 5-phosphate reductoisomerase) are important enzymes (Zhang et al. 2022) in the MEP pathway. This pathway produces the GPP (geranyl diphosphate) and GGPP (geranylgeranyl diphosphate), the precursors of monoterpenes and carotenoids, respectively (Xu et al., 2016). Therefore, there may be a possible competition in production of these compounds (Mostafa et al., 2022; Yeon et al., 2021).

Although Iranian narcissi are so popular for their strong and pleasant aroma, but no research has been carried out to compare the scent and color produced from flower tissues (i.e. the corona and perianth) of Iranian narcissus accessions. The aim of the present study was to evaluate the scent and color production separately in corona and perianth of eleven Iranian narcissus accessions, for a better understanding of the bio-physiological differences and bioactive compounds in these tissues for further use in the floriculture industry.

## 2. Materials and Methods

### 2.1. Plant material

Flower bulbs of eleven Iranian narcissus accessions were collected from natural habitats and old gardens across Iran (*N. papyraceus* and Shahla genotype from Shiraz; Shahla genotype from Kazerun; Porpar genotype and two accessions of Shahla from

Khafr; Shahla and Meskinak genotypes from Behbahan, Shahla genotype from Khusf, Shahla genotype from Juybar; Shahla genotype from Abdanan). It is worth mentioning that Shahla, Porpar, and Meskinak genotypes belong to the *N.tazetta* species (Fig. 1).

**Figure 1.**
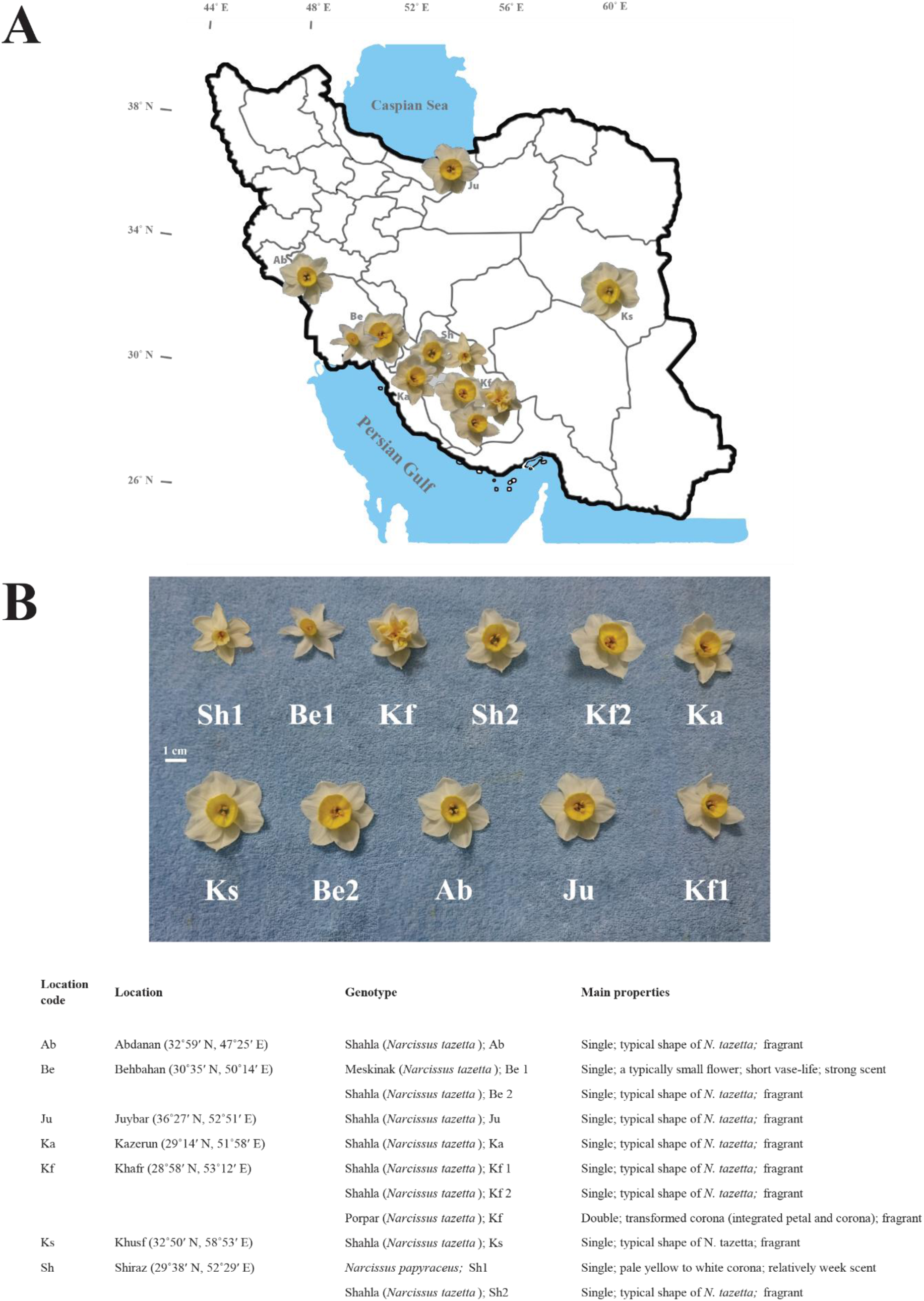
The narcissus genotypes collected from across Iran. (A) Distribution map of the narcissus accessions. (B) Flower types, geolocation of collection sites and distinct morphological characteristics.

The bulbs were cultivated in the greenhouse bed. The distance between the rows and bulbs were 20 and 18 cm, respectively. The silty-clay soil was mixed with leaf mold with a 4:1 ratio. The average ambient temperature was set as 23 ± 5°C during the growing season and natural photoperiod was used.

### 2.2. Analysis of volatile compounds

Full opened flowers were harvested at 8 am. Then, the corona and perianth were separated carefully; and one gram from each of these tissues of various genotypes were placed in two vials and sealed immediately. The vials were incubated in 80°C for 20 minutes using PAL COMBI-xt Headspace. The floral scent qualitative identification was performed by gas chromatography/mass spectrometry (Agilent Technologies, 5977B, USA) equipped with a 30 m × 0.250 mm HP-5MS column with a film thickness of 0.25 μm. The injector temperature was maintained at 280°C. The oven temperature was started at 60 °C and then raised to 210°C at 3°C/min. Thereafter, the temperature was increased to 240°C at 20°C /min and held for 8/5 min. The carrier gas was helium with a flow rate of 1 mL/min, and the electron energy was 70eV. The volatile compounds were identified by comparing their spectra with those recorded in a MS library using the standard reference database NIST. In addition, the constituents were confirmed by matching Kováts retention indices and spectra data with literatures. The integrated area of each peak relative to the total integrated peak area was used to calculate the relative percentage of volatile compounds.

### 2.3. Carotenoid measurement and colorimetry

The modified method of Hiscox and Israelstam (1978) was used for extracting total carotenoid from flower tissues (the only difference with the original method was that in the present study the tissues were soaked in DMSO for 24h in the ambient temperature). Carotenoid content was determined according to the spectrophotometry method of Wellburn (1994). Colorimetry of corona and perianth tissues was carried out by Konica Minolta colorimeter (Japan).

### 2.4. Sensory Analysis

Sensory analysis of floral tissue scent was conducted using five Shahla genotypes. Similar to the operation of preparing flowers for analyzing volatiles by GCMS, one gram of perianth and one gram of corona of each genotype were placed in two dark vials, separately; in such a way that the tissues could not be seen from outside of the vials. Each vial was identified by a code. Forty non-trained people from School of Agriculture of Shiraz University were participated in sensory analysis, with no age limitation. The tissue scent assessment was carried out using paired comparison test. Participants were asked to give score of zero or one to the tissues with lower and higher scent intensity, respectively. Thereafter, the percentages and sum of tissue scores of each genotype, and also the overall score of tissues for all genotypes were calculated.

### 2.5. Tissue micromorphology

A Scanning Electron Microscope (SEM) along with a polarizing light microscope were used to evaluate the surface morphology of perianth and corona. For Scanning Electron Microscopy, fabricating replica from both sides of mature perianth and coronas was performed using a piece of warmed dental wax (Ghanbari et al., 2019). Then, the negative duplicate was filled by apoxy resin and a hardener (with weight ratio of 2:1). The positive replica was photographed by the SEM TESCAN-Vega 3 (Brno, Czech Republic).

A polarizing light microscope (Olympus, BH41) with transmitted light, along with a smartphone camera (Sony Xperia Z5) were employed to capture images from both sides of mature perianth and corona. For this purpose, fresh mature perianths and coronas were used. Epidermal cell number were counted by ImageJ software. Tissue thickness was measured by a 0.01 mm micrometer (Asimeto, Germany). Furthermore, fresh weight of flower tissue was measured for all genotypes.

### 2.6. Primer design

The sequences of PSY, DXS, DXR genes and Actin (as internal control gene) were taken from the NCBI database and the primers were designed with AlleleID 7.5 software. Primer design was carried out according to the alignment of sequences with some *N. tazetta* RNA-seq data for GGPPs, Ocimene synthase (TPS), CCD1 and CCD4 genes. The data of these genes for narcissus is not available in the literature. Therefore, the protein sequences of these genes were taken from the databases of NCBI, InterPro Scan, Uniprot and Expasy and then their functions and domains were checked. The data of transcriptomes of perianth and corona, including accessions SRP083092, SRP083093, SRP063958, and GSE126727 were retrieved from GenBank Short Read Archive. To create contings, de novo assembly was performed using CLC Genomics Workbench 20. The BlastX was carried out by CLC. Functions and domains of proteins (i.e. the product of the contigs) was confirmed by checking on the InterProScan, NCBI, Uniprot and Expasy. The sequences with desired functions were selected. The conserved regions of the contigs and homologous sequences of other plant species were determined by Vector NTI 10 software. Then primers were designed using AlleleID 7.5 from conserved regions (Table 1). Thereafter, primer-blast was carried out on NCBI and eventually, primers were synthesized by Metobion company (Germany).

**Table 1.**
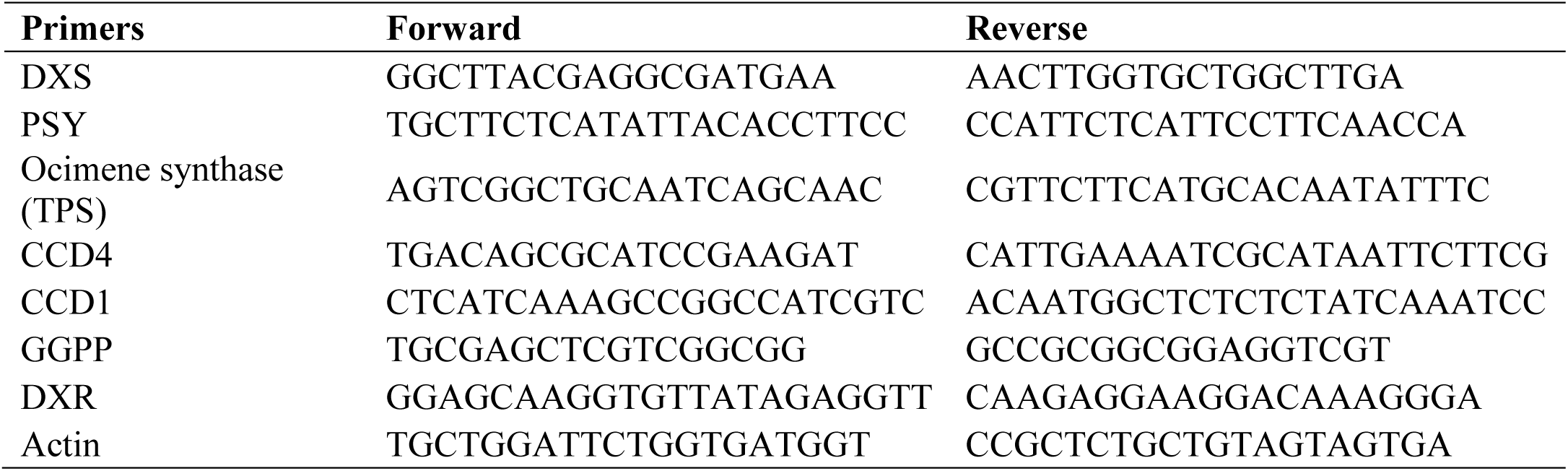
The primers used for gene expression analysis.

#### The accession numbers used

**CCD:** ACP27629.1, AJP16418.1, AJP16419.1, NP_191911.1, AAT68189.1, BBB22182.1, AZP54317.1, AAT68187.1, AAK00622.1, AYK03324.1, AYK03325.1; **GGPPs:** AAL17614.2, AA D16018.1, NP_175376.1, NP_179454.1, NP_179960.1, NP_188073.2, NP_195399.1, ADD49734.1, ADD49735.1, ABB82554.1, ABB82555.1, NP_195399.1, CAA56554.1, AAA86688.1; **Ocimene synthase:** AHI50306.1, ASN64415.1, AHI50307.1, ASN64414.1, A0A2U1KKG9|, QYM90041.1, AAN65379.1, QID05625.1, BAD91046.1, A0A1V0QSG1.

### 2.7. RNA and DNA extraction

In order to select two narcissus genotypes with the highest and lowest differences between corona and perianth in the amount of carotenoid and volatile organic compounds, Hierarchical clustering was used. For this purpose, the difference between amount of carotenoid of corona and perianth, and also the difference between the VOC relative percentages in the two tissues were calculated separately. Then, two genotypes from the clusters with the highest and lowest values of VOCs and carotenoids, were selected for gene expression analysis; i.e. Juybar & Khafr2, respectively.

Total RNA was extracted separately from 100 mg perianth and corona using Column RNA isolation kit (Dena Zist Asia, Iran). RNA quality was assessed by electrophoresis of the RNA samples on 1% agarose gel and its quantity was determined by Nano-Drop 1000 spectrophotometer (Wilmington, USA). RNA was treated with DNase I (Fermentas Thermo Fisher Scentific, USA). Then cDNA was synthesized by First Strand cDNA Synthesis kit (SinaClon, Iran). All procedures were followed according to the manufacturer’s protocols.

### 2.8. Quantitative real-time PCR

Relative real-time PCR was performed in 20 μL, including 10 μL RealQ Plus 2× Master Mix Green (High Rox; ampliqon, Denmark), 0.5 μL forward primers, 0.5 μL reverse primers, 2 μL diluted template (1:3), and 7 μL ddH_2_O using the ABI Step One (Thermo Fisher Scientific, USA) with two technical replicates. The reaction condition was as follows: an initial denaturation at 95°C for 15 min followed by 40 cycles of 95°C for 20s, Tm for 30s and 72°C for 30s. Afterwards, the temperature program 95-45-95°C with a temperature increment of 1°C every 1 min was conducted for melting-curve. The 2^−ΔΔCt^ method was used to calculate the relative gene expression (Livak and Schmittgen, 2001).

### 2.9. Experimental design

This study was conducted as a factorial test in a Completely Randomized design (CRD) with three replications for every assessment. The treatments were eleven genotypes of narcissus and 2 floral tissues (corona and perianth). Analysis of variance was performed by SPSS 21 software. Comparing treatment means was carried out using the Duncan test. Principal component analysis and Pearson’s correlation test was conducted using XLSTAT Version 2016.02.28451.

## 3. Results and discussion

### 3.1. Scent and color production

The major scent compounds detected in the whole narcissus flower were comparable with the previous researches. In this study on different flower tissues, the volatile organic compounds eucalyptol, Z and E-β-ocimene, linalool, and benzyl acetate were detected in perianth and corona of most genotypes, among which the most abundant compound was E-β-ocimene. However, this compound was 8% higher in perianth, compared with the corona tissue (Fig. 2, and 3). It has been reported that E-β-ocimene is the most abundant volatile compounds emitted from narcissus flowers (Hsin-chun chen, 2013; Ruíz-Ramón, 2014; Zarifikhosroshahi, 2021). Moreover, Knudsen (2006) and Farré-Armengol et al. (2017) reported that E-β-ocimene was present in 71% of the families of Angiosperms. The widespread distribution of such an abundant substance in flowers is assumed to have other roles than pollinator attraction (Knudsen, 2006). The second main compound found in in the corona and perianth of the *N. tazetta* genotypes was benzyl acetate and linalool, respectively. It was in accordance with previous studies which reported that the major benzenoid in *N. papyraceus* and *N. tazetta* var. *chinensis* is benzyl acetate (Dobson et al. 1997; Song et al., 2007, Hsin-chun chen 2013, Ruíz-Ramón, 2014; Zarifikhosroshahi, 2021).

**Figure 2.**
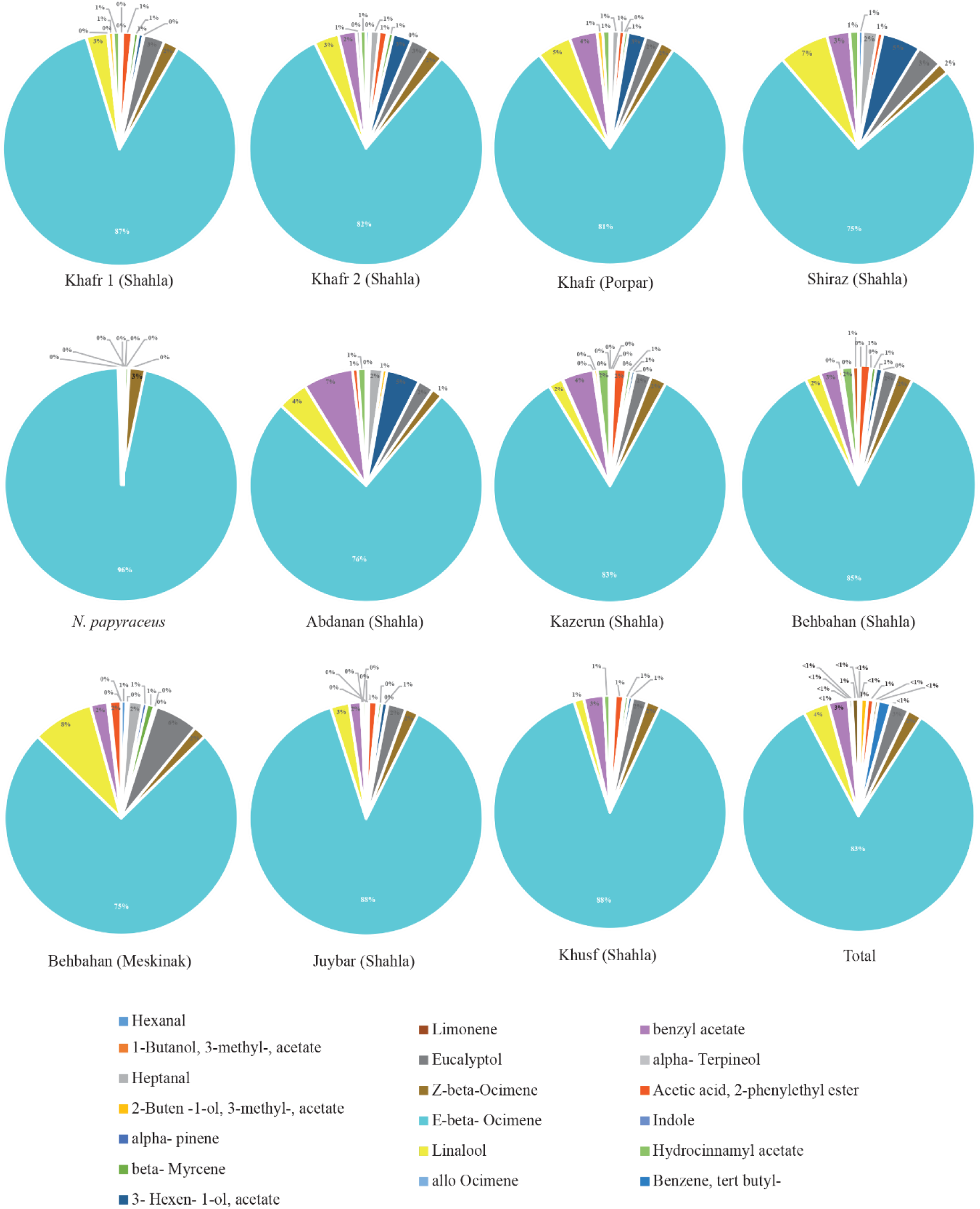
Percentage of volatile organic compounds measured in the perianth tissue of Iranian narcissus genotypes (*N. tazetta*). The zero values show the relative percentages of compounds less than 1%.

However, it was revealed that the flower tissues did not emit the same VOCs in all genotypes. Alpha-terpineol, acetic acid, 2-phenylethyl ester, α-pinene, β-myrcene, and limonene were detected in corona of all genotypes of *N. tazetta*, but not in perianth of all genotypes (Fig. 2, and 3). Alpha-terpineol was detected only in perianth of two genotypes, while 2-phenylethyl ester, β-myrcene, limonene, and α-pinene were identified in the majority of narcissus genotypes. Furthermore, indole was identified only in corona of Shahla genotypes in Kazerun and Behbahan (Fig. 3). The findings of this study demonstrates that the differences between quality and relative quantity of flower VOCs, detected in perianth and corona, is affected by genotypes.

**Figure 3.**
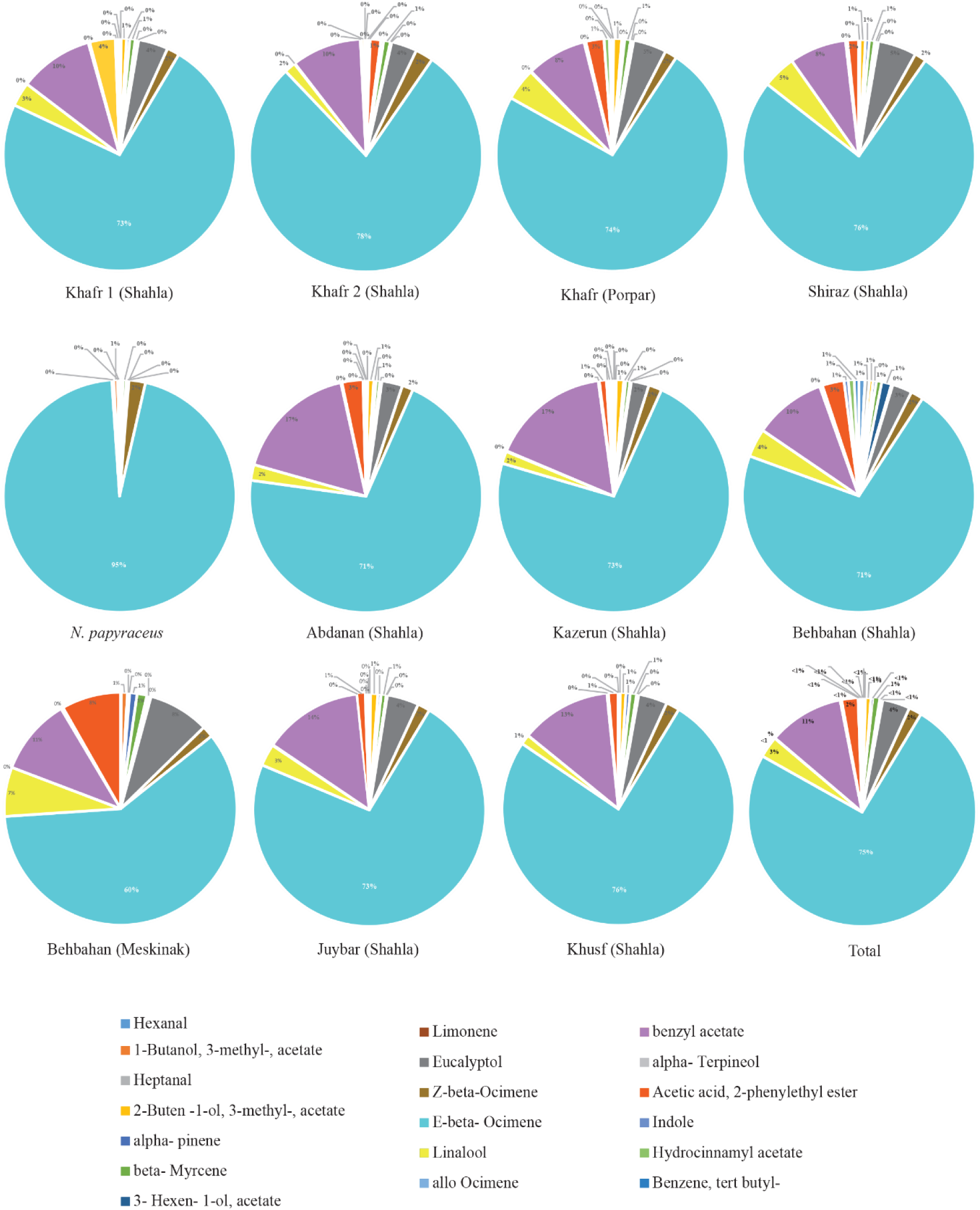
Percentage of VOCs detected in the corona tissue of Iranian *N. tazetta*. The zero values show the relative percentages of constituents less than 1%.

The number of identified compounds by GC-MS varied among different tissues and genotypes of narcissus (Fig. 2, and 3). The least number of constituents among the genotypes was seen in *N. papyraceus*. Furthermore, α-pinene, limonene, eucalyptol, α-terpineol, and acetic acid, 2-phenylethyl ester were not detected in the tissues of *N. papyraceus*. In contrast, the relative percentage of E-β-ocimene measured in its corona and perianth tissues was higher than other genotypes. Compared to the other genotypes, the corona and perianths of Meskinak had a higher relative percentage of β-myrcene, eucalyptol, linalool, acetic acid, 2-phenylethyl ester, but less E-β-ocimene (Fig. 2, and 3). Ruiz-Ramon et al. (2014) have reported different percentages and types of volatile compounds in single and double varieties of *N. tazetta*. Also, previous researches have shown that floral volatile compounds may vary among closely related species (Dobson et al., 1997; Knudsen, 2006), or even between plants and between populations (Dudareva, 2006).

The amount of carotenoids observed in the corona tissue of all narcissus genotypes was higher than perianth. Among all of the genotypes, Meskinak had more carotenoid in both tissues, while the least carotenoid of corona was observed in *N. papyraceus* (Fig. 4). As shown before, Valadon and Mummery (1968) indicated that variations exist in the type and amount of carotenoids in different parts of narcissus flowers, depending on genotypes, maturity level and growing conditions. The findings of the current study are consistent with the previous report.

**Figure 4.**
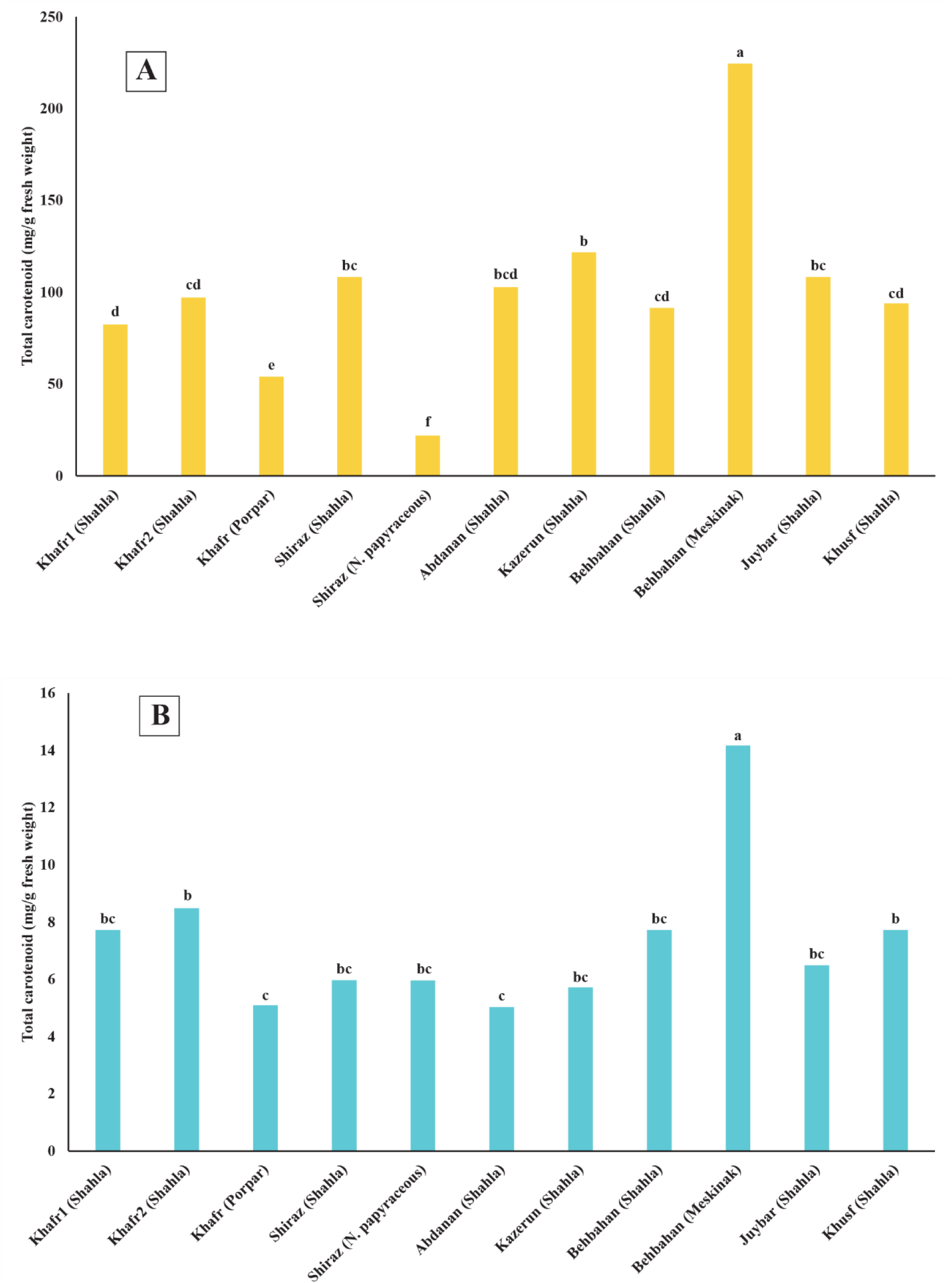
Total carotenoid measured in the corona (A) and perianths (B) of Iranian narcissus genotypes (*N. tazetta*). Different letters indicate significant differences of various genotypes (*p* ≤ 0.01).

In the corona of narcissus flowers, the amount of carotenoids had a significantly negative correlation with E-β-ocimene relative percentage, but a significantly positive correlation with acetic acid, 2-phenylethyl ester and monoterpenes α-pinene, eucalyptol and β-myrcene (Fig. 5). Also, in corona of all genotypes there were significant correlations between a* and L* (CIELab color space) and the two compounds of E-β-ocimene and benzyl acetate. Furthermore, in perianth of narcissus flowers, the carotenoids had significant positive correlations with α-pinene, eucalyptol, β-myrcene and limonene (Fig. 5). The relationship among color and amount of total carotenoids and the expression level of genes related to color is shown in Figure 6. The factors a* and b* had positive correlations with total carotenoids and PSY gene expression level. Moreover, the parameter L* which shows lightness of color, had a positive correlation with the expression of CDD4 gene, and a negative correlation with total carotenoids (Fig. 6). Li et al. (2015) identified ten carotenoids in various cultivars of the narcissus flowers. They indicated that all-*trans*-violaxanthin and total carotenoid were the main pigments affecting the color of narcissus flowers where L* and b* parameters were increased, but a* was decreased. Findings of the present study differed with the reports of Li et al. (2015), as the total carotenoid of Iranian narcissi here had positive correlation with a* and b*, and negative correlation with L*. Considering the strong correlations between the amount of carotenoids and color values (L* a* b*), the significant relationship between color values and E-β-ocimene is also reasonable.

**Fig 5.**
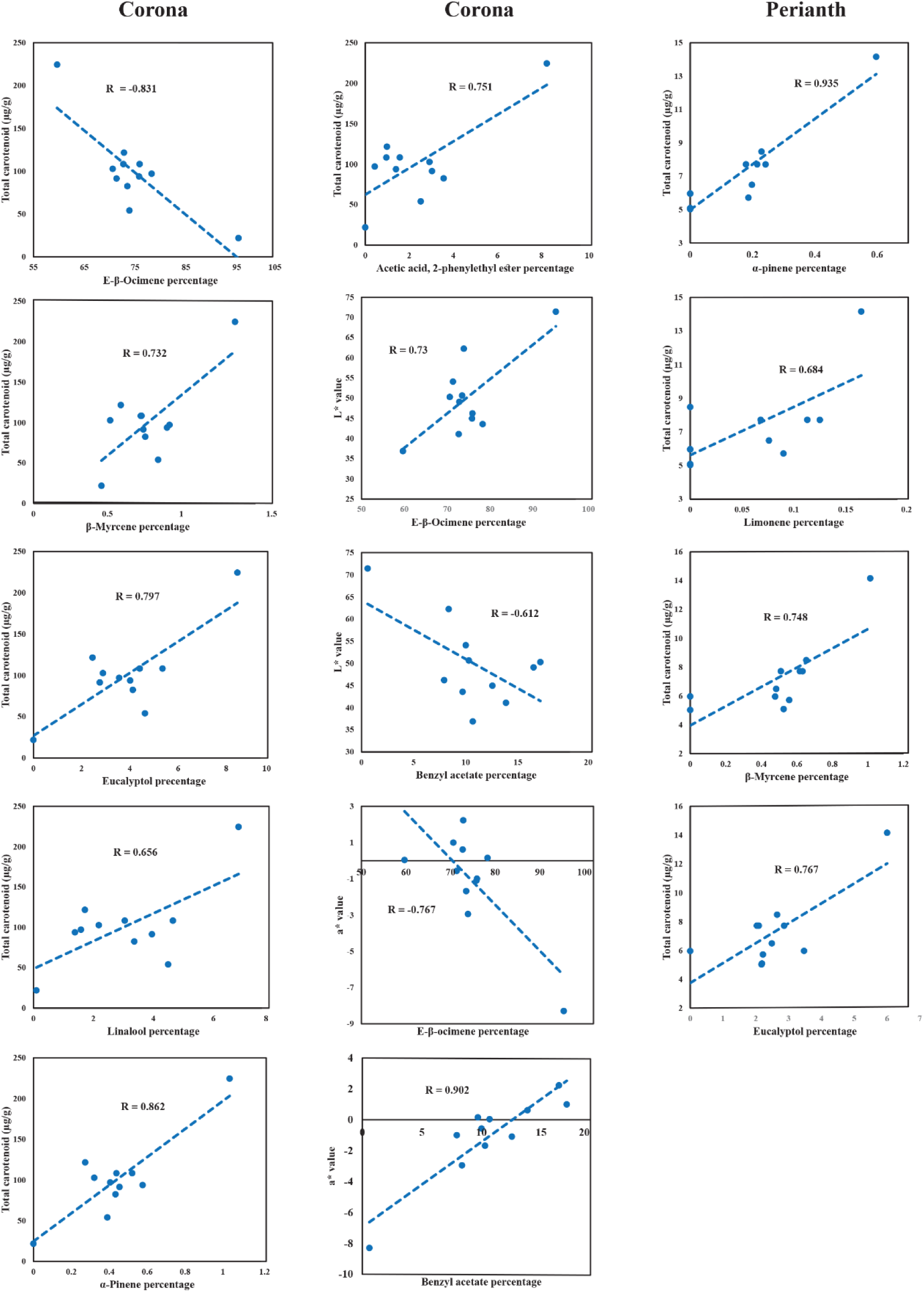
Significant correlations detected in the corona and perianth of Iranian narcissus flowers, between total carotenoid and some VOCs, and also between color values (a* and L* of CIELab color system) and volatile organic compounds.

**Fig 6.**
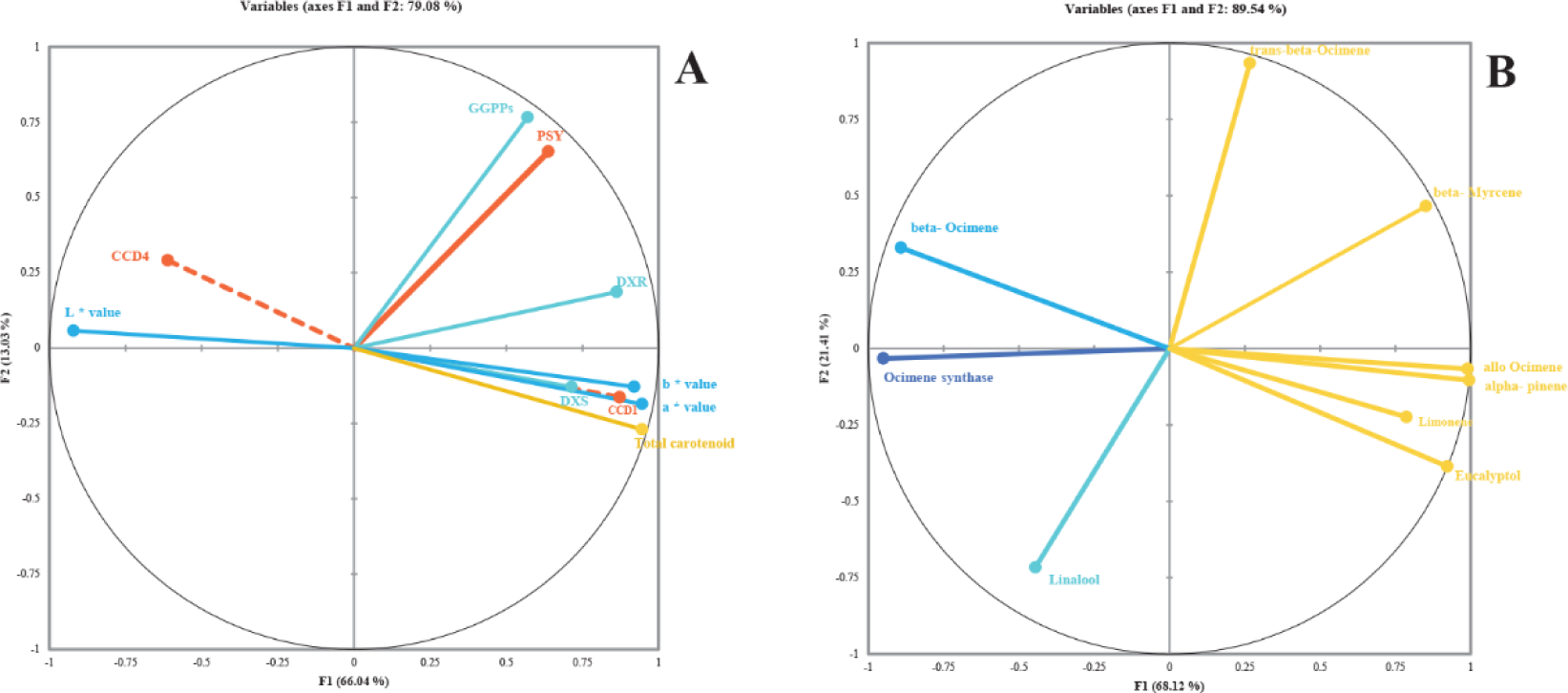
Principal component analysis for A) total carotenoid, color values and the expression of the genes associated with carotenoid production and B) volatile organic compounds production and ocimene synthase gene expression in the Iranian narcissus flowers.

It is has been known that TPSs produce several compounds (Dudareva, 2006). However, principal component analysis of the present study showed that the variations observed in the expression of terpene synthase was consistent with E-β-ocimene emission, as it is the dominant monoterpenoid VOC of narcissus flower. It should be noted that such pattern was not observed for the other monoterpenes (Fig. 6b). This shows that TPS expression of narcissus flowers is mainly related to the E-β-ocimene synthesis. In a previous study, Farré-Armengol et al. (2017) have shown that the ocimene constituent is the main product (95%) of ocimene synthase gene, though a small amount of other monoterpens like myrcene are also produced. Yet, further studies are required for a better understanding of ocimene synthesis in the narcissus flower.

### 3.2. Flower morphology & Scent perception

The fresh weight of perianth was higher than corona tissue. Shahla genotype showed 47% increase in perianth’s fresh weight, compared to the other genotypes. However, the thickness of the corona was significantly greater (48%) than perianth (Fig. 7). Except for the Porpar genotype which has a transformed corona, generally perianth had more fresh weight than corona. Furthermore, in all genotypes, except in Khusf (Shahla), Behbahan (Meskinak), and Shiraz (*N. papyraceus*), corona tissue was significantly thicker than perianth (Fig. 7). It is commonly assumed that thicker perianth prolong the flower’s vase life, but decrease its scent production. According to the results of the sensory analysis (Fig. 8), coronas had stronger fragrance than the perianth. However, in the rose flower Bergougnoux (2007) showed that scented and non-scented varieties of roses did not have any significant differences in their petal thickness. In accordance with the previous finding, in the present study no considerable correlation was observed between scent perception neither with tissue fresh weight, nor with tissue thickness (data not shown).

**Fig 7.**
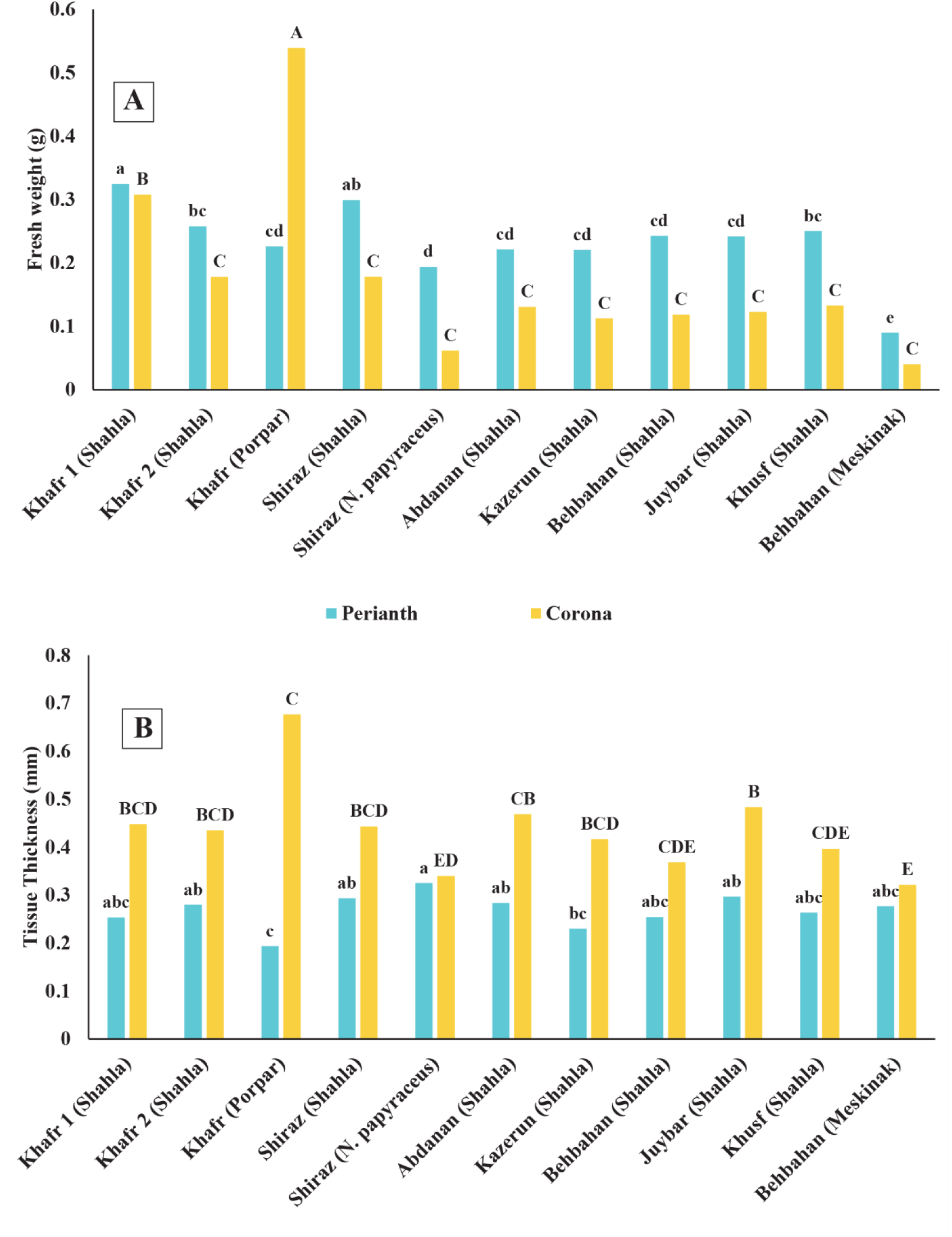
Perianth and corona fresh weight (A) and thickness (B) of Iranian narcissus genotypes (*N. tazetta*). Different small and capital letters indicate significant differences among the perianth, and also among the coronas of various genotypes, respectively (*p* ≤ 0.01). T-bars show standard deviation.

**Fig 8.**
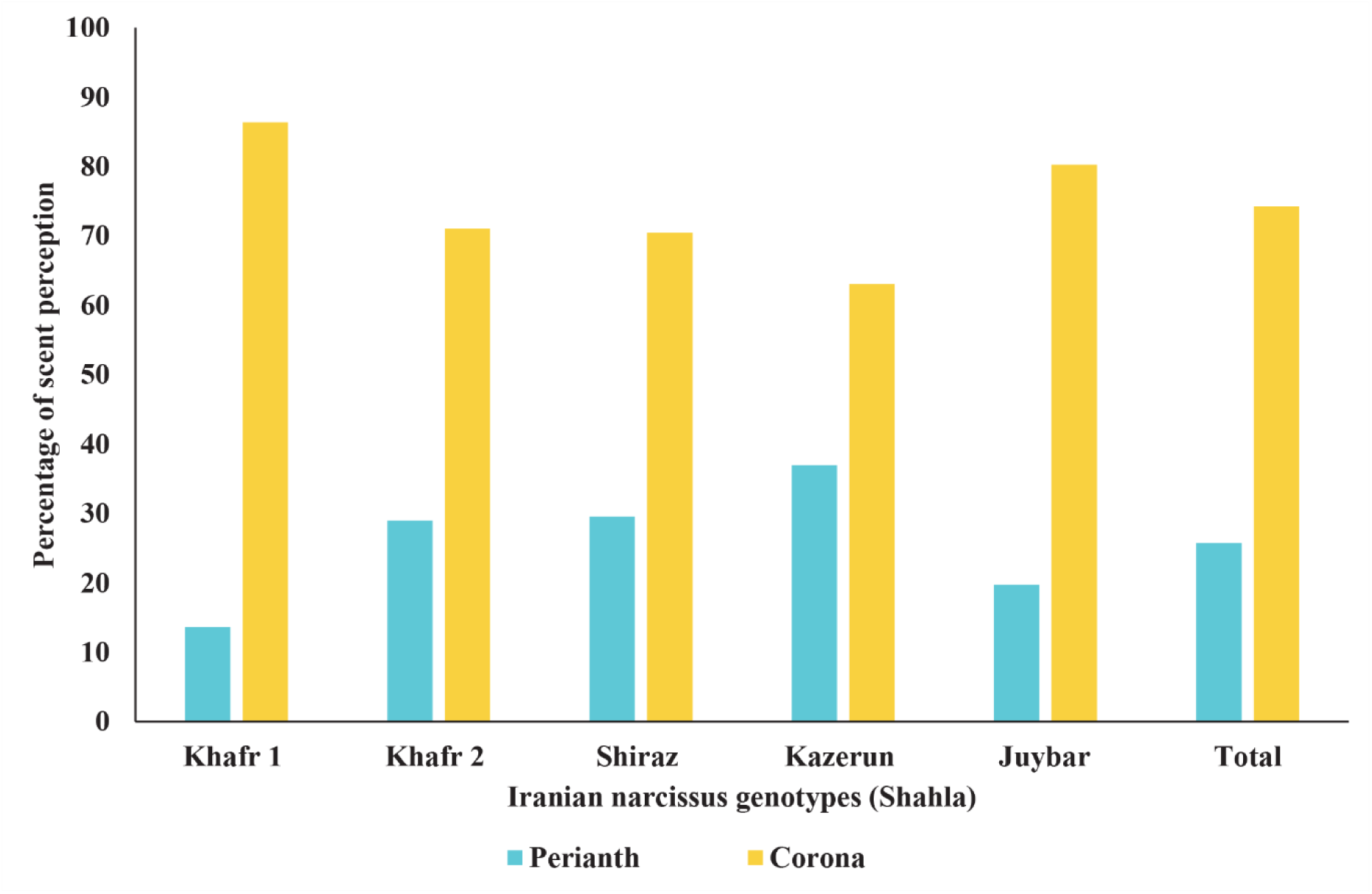
Comparatively scent perception from perianth and corona of 5 Iranian narcissus genotypes (Shahla).

Results of sensory analysis indicated that the corona had a bigger scent perception score, 2.9 times higher than perianth (74.47% vs 25.52%), which reveals that the participants in the sensory analysis had felt 48% more intensity of scent from the corona, than from the perianth (Fig. 8). Despite the high percentage of E-β-ocimene in narcissus flower tissues, it implies that other VOCs such as benzyl acetate, α-terpineol, acetic acid, 2-phenylethyl ester, α-pinene, β-myrcene, and limonene possibly have a greater role in fragrance perception by humans. This is because in the equal weight of corona and perianth tissues, the percentage of the latter compounds were either higher in the volatile mixture of the corona than perianth, or these compounds were detected just in the corona. Findings of the present study suggest that quality and relative quantity of VOCs were different in perianth and corona, while genotypes had considerable effects on these differences as well. In addition, it is revealed that corona emits higher levels of scent compared to the perianth. Moreover, it is also shown that narcissus corona with different shapes and colors, has probably more important roles in flower aesthetics, attraction, signaling and repellence of insects, than known before.

### 3.3. Flower microscope analysis

Polarized light microscopy showed that the abaxial and adaxial surfaces of the perianth and corona were covered with shiny white and yellow conical cells, respectively. The cells had almost the same size and density on both sides of perianth and corona, but not in all of narcissus genotypes (Fig. 9). Interestingly, scanning electron microscope revealed more details about the conical cells’ surface. The epidermal cells of perianth and corona had cuticular striations. To the best of our knowledge, this is the first report showing the similarity of the conical cells on double side of perianth and corona, and also the first hint for the presence of cuticular striations on epidermal cells of these tissues in the narcissus flower. Previous studies have demonstrated that conical cells are the place where pigments are produced and scent is released (Cavallini-Speisser, 2021; Yeon, 2021). These cells also increase tissue color intensity and as a consequence contribute to its shiny appearance (Cavallini-Speisser, 2021). Comparatively, in the rose flowers, scent volatiles are produced and emitted by both the adaxial and abaxial epidermal petal layers. It was found that the shape of conical cells reflects the light and guides the pollinating insects to the flowers. Those flowers with flat epidermal cells attract less insects, because they reflect less light (Bergougnoux, 2007). The findings of the current study which show that conical cells of perianth and corona are shiny are in line with previous reports.

**Fig 9.**
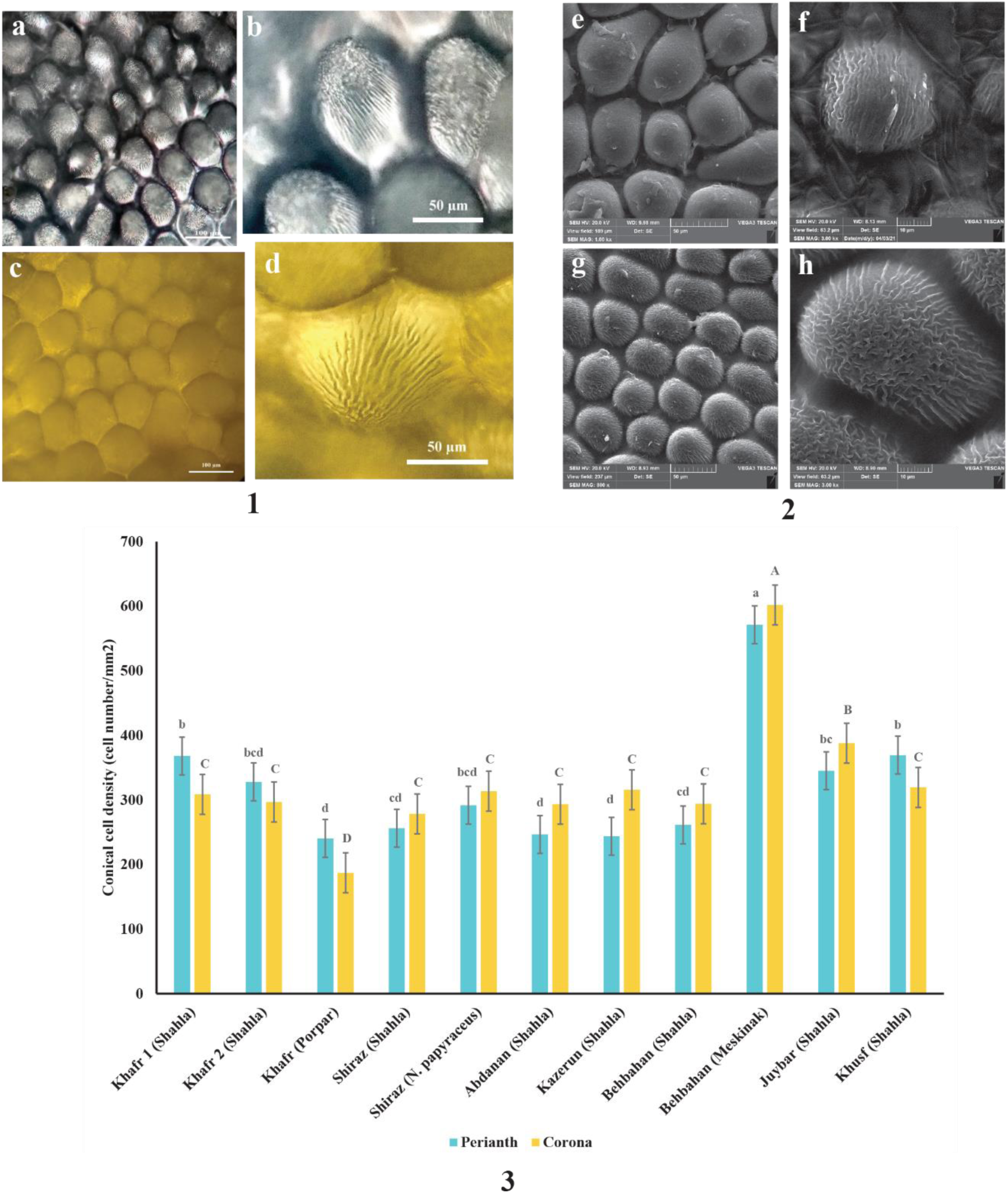
Conical cells of perianth and corona of Iranian narcissus genotypes. Images were captured by polarized light microscopy with transmitted light from using fresh unstained tissues (1); and scanning electron microscope (SEM) (2). a, b, e, and f: conical cells of perianth; c, d, g, and h: conical cells of corona. Part (3) shows the conical cell density in perianth and corona of Iranian narcissus genotypes (*N. tazetta*). Different letters indicate significant differences among the perianth, and corona of various genotypes, respectively (*p* ≤ 0.01; T-bars show standard deviation).

### 3.4. Gene expression

While in the two evaluated genotypes, the expression of DXS, DXR, GGPPs, PSY, and CCD1 genes were higher in the corona than perianth, the expression of ocimene synthase and CCD4 gene was higher in the perianth. In general, the expression of CCD4 gene was higher than CCD1, in both tissues (Fig. 10). Ren et al. (2017) showed that among the carotenoid degradation genes, NCED, CCD7, and CCD8 had extremely low expression in *Narcissus* perianth. In the current study, although CCD1 and CCD4 had the major role in carotenoid degradation, expression of CCD4 is much higher than CCD1. This observation is consistent with the reports of Ren et al. (2017). In the Iranian narcissus flowers, CCD4 is the main responsible gene for carotenoid degradation.

**Fig 10.**
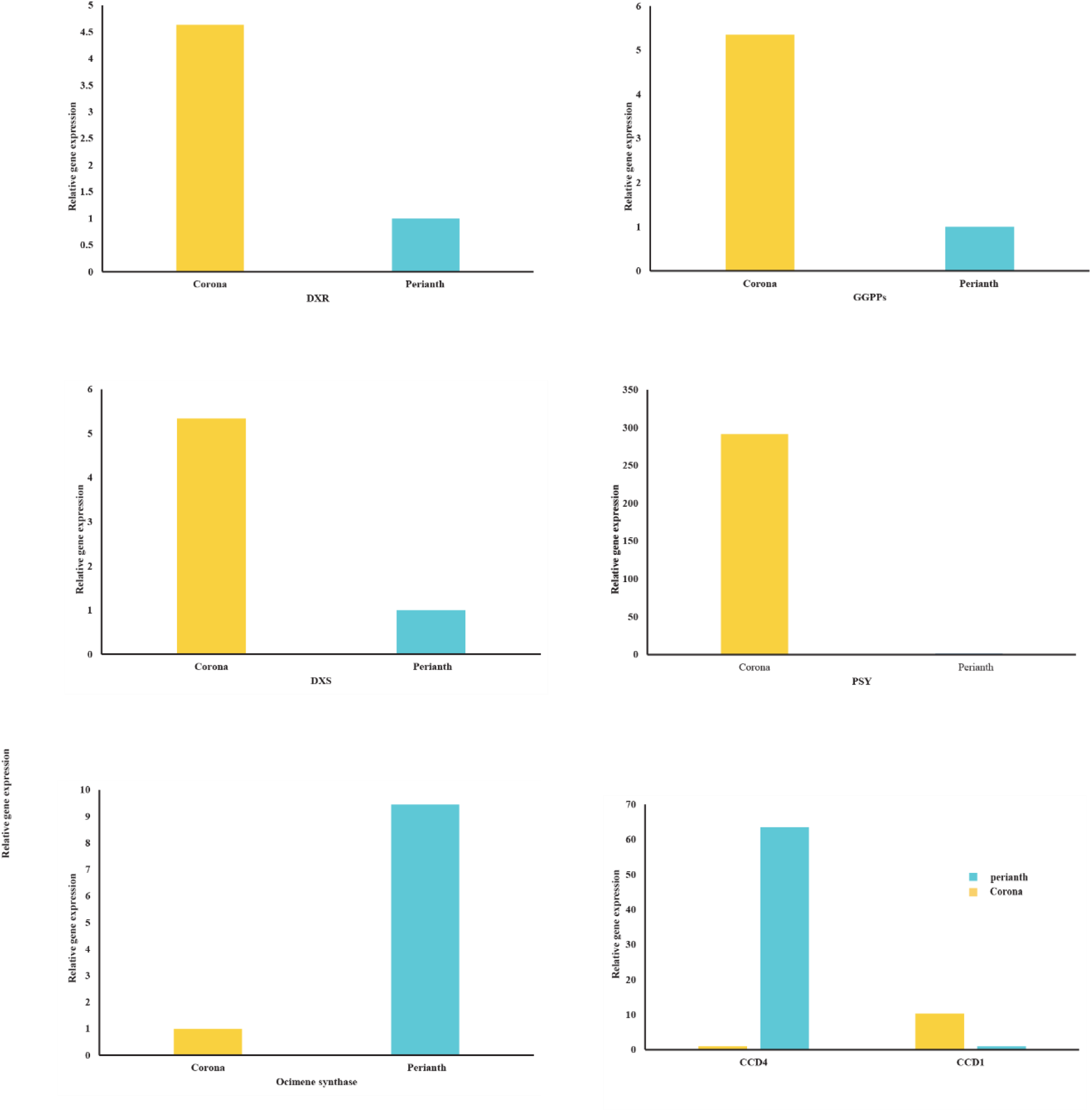
The relative gene expressions associated with color and scent in perianth and corona of Iranian narcissus genotypes (Shahla).

Although there was significant negative correlation between the measured values of E-β-ocimene and carotenoid, the correlations between the expression level of ocimene synthase and PSY, and also between expression of these two genes with DXR was positive (R = 0.82, and R=0.56, respectively) (Table 2). Furthermore, the variations of DXR, DXS, GGPPs, and PSY genes expression were almost in the same directions (Fig. 6-a) (Table 2).

**Table 2.** Pearson correlation between gene expression levels related to color and scent in corona of Iranian narcissus genotypes (Shahla).

| Variables | CCD1 | PSY | DXS | DXR | GGPP | CCD4 |
| --- | --- | --- | --- | --- | --- | --- |
| CCD1 | **1** |  |  |  |  |  |
| PSY | 0.034 | **1** |  |  |  |  |
| DXS | **0.420** | 0.193 | **1** |  |  |  |
| DXR | **0.453** | **0.616** | **0.559** | **1** |  |  |
| GGPP | 0.116 | -0.003 | 0.018 | **0.422** | **1** |  |
| CCD4 | -0.178 | -0.100 | -0.152 | -0.214 | -0.067 | **1** |
| Ocimene synthase | 0.293 | **0.788** | **0.445** | **0.823** | 0.194 | -0.131 |

*Values in bold are different from 0 with a significance level $\alpha=0.05$*
Values in bold indicate the significance correlations ( $p < 0.05$ ).
CCD 1 & 4: carotenoid cleavage deoxygenases 1 &4
DXS: 1-deoxy-D-xylulose-5-phosphate synthase
DXR: 1-Deoxy-D-xylulose 5-phosphate reductoisomerase
GGPPs: geranylgeranyl diphosphate
PSY: Phytoene synthase

It is well known that carotenoids and monoterpenes are both produced by the MEP pathway in plastids and have similar precursors (Farré-Armengol et al., 2017), so there is a competition in their production (Mostafa, 2022; Yeon, 2021). DXS and DXR are two enzymes with rate-limiting roles in the early stages of the MEP pathway (Xu et al., 2016; Zhang et al., 2022), which play the same role both in synthesis of carotenoids and monoterpenes. In the current study, higher expression levels of DXR and DXS in the corona, is because both carotenoid and volatile monoterpenes were produced in greater amounts in the corona, than inside perianth. Moreover, the orange corona had higher expression of GGPPs than perianth. Geranylgeranyl Diphosphate (GGPP), the precursor of carotenoids, is produced by GGPPs (Xu, 2016; Yang et al., 2021). Yang et al., (2021) reported that there is a competition between carotenoid and terpene backbone biosynthetic branches for GGPP in the narcissus flower tissues. Zhang et al. (2022) indicated that DXR and PSY had higher expression levels in the perianth of yellow narcissus compared with the white type (Zhang et al., 2022). The similar trend of expressions of DXS, DXR, GGPPs, and PSY, is due to the role of these genes in MEP pathway and carotenoid biosynthesis.

## 5. Conclusion

The color and scent analysis of various Iranian narcissus flowers revealed differences not only among the accessions, but also among the corona and perianth of each genotype. The E-β-ocimene was the dominant volatile organic compound identified in both tissues of all genotypes. Meskinak had the highest amount of carotenoid and also different number of VOCs compared to other genotypes. Moreover, there was a linkage between color and scent in narcissus flowers which suggest a similar pathway and therefore a competition for production of monoterpenes and carotenoids in corona. Volatile compound percentages, color, scent emission, fresh weight and thickness of tissue were different in the perianth and corona of narcissus flower, while the type of volatile organic compounds and the surface coverage (with epidermal conical cells) were similar. Although both perianth and corona play important roles in attracting pollinators and also humans, it seems that corona has a stronger effect in the Iranian narcissus flowers. The findings of this research for the first time clearly illustrates the distinguished role of corona in greater production of color and scent in Iranian narcissus flowers and can be used in the floriculture industry and aromatherapy.

